# Utilizing flow cytometry sorting signal width to enrich for cells positive to endogenous gene integration of fluorescent proteins

**DOI:** 10.1101/2023.03.21.533670

**Authors:** Gabriel P. Faber, Hagit Hauschner, Mohammad Khaled Atrash, Liat Bilinsky, Yaron Shav-Tal

## Abstract

Endogenous gene knock-in using CRIPSR is becoming the standard for fluorescent tagging of endogenous proteins. Some protocols, particularly those that utilize insert cassettes that carry a fluorescent protein tag, can yield many types of cells with off-target insertions that have diffuse fluorescent signal throughout the whole cell in addition to scarce cells with on-target gene insertions that show the correct sub-cellular localization of the tagged protein. As such, when searching for cells with on-target integration using flow cytometry, the off-target fluorescent cells yield a high percentage of false positives. Here, we show that by changing the gating used to select for fluorescence during flow cytometry sorting, namely utilizing the width of the signal as opposed to the area, we can highly enrich for positively integrated cells. Reproducible gates were created to select for even minuscule percentages of correct subcellular signal, and these parameters were validated by fluorescence microscopy. This method is a powerful tool to rapidly enhance the generation of cell-lines with correctly integrated gene knock-ins encoding endogenous fluorescent proteins.

## 1 INTRODUCTION

Targeted integration of a sequence to tag an endogenous gene, also termed knock-in (KI), is rapidly becoming the standard approach in the generation of endogenous fluorescent protein (FP) tagging. Many different protocols using CRISPR Cas9 or Cas12a have been developed [1-3]. One particular method [4] uses an integration cassette that is synthesized using PCR, which contains not only the homology arms and the coding region of a fluorescent protein, but also the guide RNA (gRNA) which is transcribed in vivo. Transfection of the PCR cassette along with Cas12a, results in integration at the 3’ of the coding gene locus and C-terminal tagging of the endogenous protein, using homologous recombination (HR) following DNA cleavage, which is active in late S/G2 phase during the mammalian cell cycle [5]. An accompanying web portal (http://www.pcr-tagging.com/) allows for generation of primers used for the homology arms with ease, and also directs which Cas12a to use based on varying PAM sequences.

Correct on-target gene integration will yield cells expressing the fluorescently tagged proteins in the correct subcellular localization. However, a major obstacle encountered is the presence of a high number of cells with off-target integrations exhibiting a fluorescent signal that is diffuse throughout the cell. For instance, on-target tagging of a nuclear protein can be very rare within a population of many off-target cells with prominent cytoplasmic signal. While the nature of this off-target signal is not entirely clear, it is sufficient to make selecting positively integrated cells a challenge. Standard flow cytometers measure the fluorescence intensity of the cell but cannot determine the exact cellular localization of the fluorescent protein. Moreover, traditional methods of selection and sorting of cells expressing fluorescent proteins with flow cytometry are not equipped to differentiate between on-target integration in the nucleus and diffuse signal of the same intensity. To address this issue, we generated gates for flow cytometry that utilize the width of the fluorescent signal, as opposed to the area which is generally employed. Because nuclear signal is confined to a much tighter spatial area, it yields a completely discernable population, which normally would appear identical to the cytoplasmic signal if looking at overall intensity. While signal width as a possible fluorescent localization indicator has been described [6], here we demonstrate the use of a new gating protocol in the pipeline of endogenous gene knock-in, to swiftly enrich for positive cells in the population, and expedite cell line generation. This method enables the distinguishing between cytoplasmic and nuclear fluorescent protein expression by leveraging of the difference in the width of the fluorescent signal, thus enabling significant positive on-target enrichment of the sorted populations.

## 2 MATERIALS AND METHODS

### 2.1 Tissue culture and transfections

Human U2OS cells were maintained in low glucose DMEM (Biological Industries, Beit-Haemek, Israel), supplemented with 10% fetal bovine serum (HyClone Laboratories, Logan, UT), 1% Glutamine (Biological Industries, Beit Haemek, Israel), and 1% penicillin–streptomycin solution (Biological Industries). Cells were grown at 37°C in 5% CO_2_.

For the knock-in of the mClover fluorescent protein into the C-terminus of the endogenous ALY/REF gene, we used the Mammalian PCR-tagging protocol for tagging the C-termini of genes [4]. The pMaCTag-P26 plasmid containing the mClover coding region along with a mini-AID was used (Addgene #120037). The online oligo design tool www.pcr-tagging.com was used for searching PAM sites and to design the forward (M1) and reverse (M2) tagging oligos specific for enAsCas12a (Addgene #107942). M1 and M2 oligos, targeting ALY/REF, were obtained from IDT.

M1:

CAGCTGGGGGCCTGGTCAAAGCCGCAGTGGGGAGCAGGCCGCCTGTGAATGCAAGCCTCTTTCTCCTCTGTGTTTCAGATGGACACCAGTTCAGGTGGAGGAGGTAGTG

M2:

AGGGAGCAAGAGGAGACGCCTGGGTCCTGTTCCGCACGCGGATTTGCTGGTCTGTAAAAAACTGTTTAACTGGTGTCCATCATCTACAAGAGTAGAAATTAGCTAGCTGCATCGGTACC

The cassette was amplified with M1 and M2, using pMaCTag-P26 as a template. To generate U2OS cells expressing endogenously tagged ALY/REF-mClover cells, ∼10^6^ cells were seeded 24 h before transfection. Plasmids containing enAsCas12a and the mClover PCR cassette were electroporated (Bio-Rad) into the cells. Cells were immediately treated with XL413, an S-phase inhibitor, and rinsed with fresh medium after 24 h to increase HDR efficiency in cells [7]. Two days later, cells were treated to Puromycin (1 μg/ml; Invivogen, San Diego, CA) for 7-14 days to select for cells with the PCR cassette insert before being prepared for sorting.

### 2.2 Sorting method

U2OS cells that underwent transfection and selection as described above were sorted using a BD FACSAria™ III sorter (BD Biosciences, San Jose, CA, United States). Parameters that were selected to acquire and record were: area, height and width (A, H, W) of the forward scatter (FSC), side scatter (SSC) and GFP (520/50) signals. At least 10,000 cells in the live cells gate were recorded. The gating strategy is described in the result section. 20,000 cells were collected and seeded for continuous growing and some (∼4000 cells) were seeded on a slide for microscopy analysis. Post-acquisition analysis was done using FlowJo software (ver. 10.8) (Ashland, OR, United states).

### 2.3 Microscopy analysis

Wide-field fluorescence images were obtained using the CellSens system based on an Olympus IX83 fully motorized inverted microscope (60× UPlanXApo objective, 1.42 NA) fitted with a Prime BSI sCMOS camera (Teledyne) driven by the CellSens software.

### 2.4 Statistical analysis

To determine significance of change in number of on target cells after sorting, a Chi-square test for independence was performed. Pearson’s Chi-squared test with Yates’ continuity correction:

*X-squared* = 261.63, *df* = 1, *p-value* < 0.00000000000000022.

## 3 RESULTS

### 3.1 Establishing a gating strategy to identify nuclear versus cytoplasmic protein localization

To begin, we surmised that cytoplasmic versus nuclear signal would show different distribution when accounting for their spatial confines. To this end, we chose distinct nuclear and cytoplasmic localized proteins, and observed them using flow cytometry by a number of metrics. To delineate signal that was restricted to the nucleus, the nuclear speckle protein 9G8 tagged with GFP was used (Figure 1A) [8, 9]. IGF2BP3 tagged with GFP was used to identify cytoplasmic signal (Figure 1B) [10]. Besides applying the standard forward and side scattering to select for singlet cells, we also measured the fluorescent signal along multiple dimensions (Figure 1A,B). In a BD FACSAriaIII sorter, a single fluorescence pulse has a normal distribution correlating to the time the cell crosses the laser beam. The pulse begins with a low intensity when a cell enters the laser beam, reaches its maximum intensity when the cell is in the middle of the beam and as it leaves the beam the pulse trails off. This pulse distribution is represented in the following parameters: the area below the graph (A), the maximum height (H), and the width which is correlated to the time of the pulse (W). Both the nuclear and cytoplasmic proteins expressed similar levels of fluorescence, and the area of the GFP signal (GFP-A) for each protein overlapped considerably (over 50%), each presenting a range between 1×10^3^ and 1×10^4^ in total intensity. It was almost impossible to discern between cells expressing each of the two proteins using this metric alone. However, when examining the distribution of the width of the signal (GFP-W), the cells showed two distinct populations that were easily distinguishable. When plotting the cells along an area versus width plot, each cell line occupied a unique spot on the graph (Figure 1A,B). The cells expressing cytoplasmic protein were situated much higher on the width axis, ranging from about 150k-200k, while the cells expressing nuclear protein were firmly below, with a width of about 75k-125k. This was seen clearly when plotting both cell lines along either an area or width axis. While the fluorescent area curves showed considerable overlap, the width curves showed two distinct peaks, with the nuclear population displaying a peak at 1×10^5^, while the cytoplasmic peak sat far to the right at 1.75×10^5^ (Figure 1C). This simple shift in data presentation was able to amply differentiate between cytoplasmic and nuclear fluorescent localized signal, which was previously unattainable.

**FIGURE 1.**
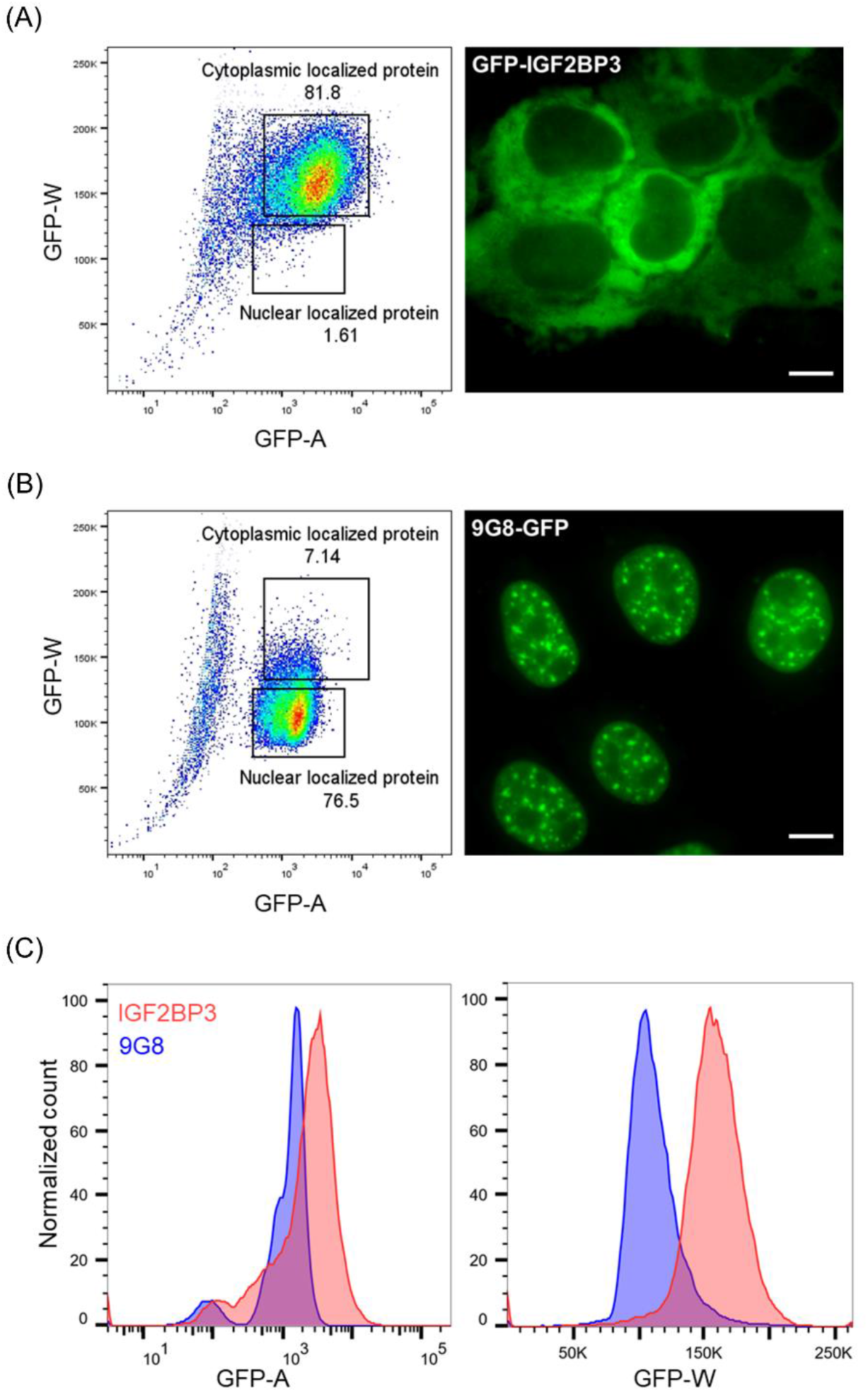
Cytometey gating by width yields distict populations. (A) GFP-A versus GFP-W distribution of U2OS stable cell lines expressing GFP-IGF2BP3. Fluorescent image shows signal demarcating the cytoplasm. (B) GFP-A versus GFP-W distribution of U20S stable cell lines expressing 9G8-GFP. Fluorescent image shows signal demarcating the nucleus. Bars, 10 μm. (C) Graphs depicting the GFP-A or GFP-W of the two proteins along an intensity axis, where they overlap considerably when looking at area of signal, but show two distict peaks when looking at the width.

### 3.2 Utilizing new sorting gates on endogenous gene integrations

In the course of studying mRNA export from the nucleus and the tagging of endogenous genes [11], we sought to tag the mRNA export factor ALY/REF (also termed THOC4) [12, 13]. This protein is known to reside in the nucleus. U2OS cells underwent transfection for endogenous gene KI, in this case with the mClover fluorescent protein (Ex:506, Em:518) using a PCR insert cassette targeting the ALY/REF gene. A few days following transfection, cells were first sorted by GFP-A in order to select for positive cells, as is traditionally done, and were imaged. The signal covered an extremely wide range of intensities, from less than 10^2^ to over 10^4^ in total mClover signal. This owed to the highly varied mClover expression in the population, which probably designates different off-target integrations and different number of copies of cassette present. In the thousands of cells that were selected in the GFP channel, very few positively-integrated cells were observed under the microscope, comprising only about 2% of the total cells (Figure 2A). This demonstrates that using this population of mixed cells for single-cell sorting would be a futile endeavor. Therefore, the same cells were taken post-transfection and examined with the new gating protocol. After transfection, the positive integrated cells, as indicated by tighter width of signal, were only about 3% of total cells, while over 75% of the population showed green signal in the cytoplasm (Figure 2B). This not only corroborated our observations under the microscope, but reiterated how many off-target cells would be collected if sorting only by GFP-A. After implementing the new gates into the sorting protocol, the post-sort population was nearly entirely enriched with on-target integrated cells (Figure 2B). While singular cells with diffuse signal were detected, after sorting for cells with positive gene integration of the nuclear protein, the number of nuclear cells exceeded 80%, significantly exceeding the prior percentage (p<0.0001). After gating, it was possible to split the GFP-A into nuclear and cytoplasmic components. Indeed, the nuclear mClover expressing cells and the cytoplasmic diffuse cells presented as two distinct populations that were easily discernable by the width of signal, although they overlapped almost entirely in GFP-A (Figure 2C). Because the same population of cells was split into their nuclear and cytoplasmic localized fractions, we could determine the GFP-W signal cutoff point at precisely 130k.

**FIGURE 2.**
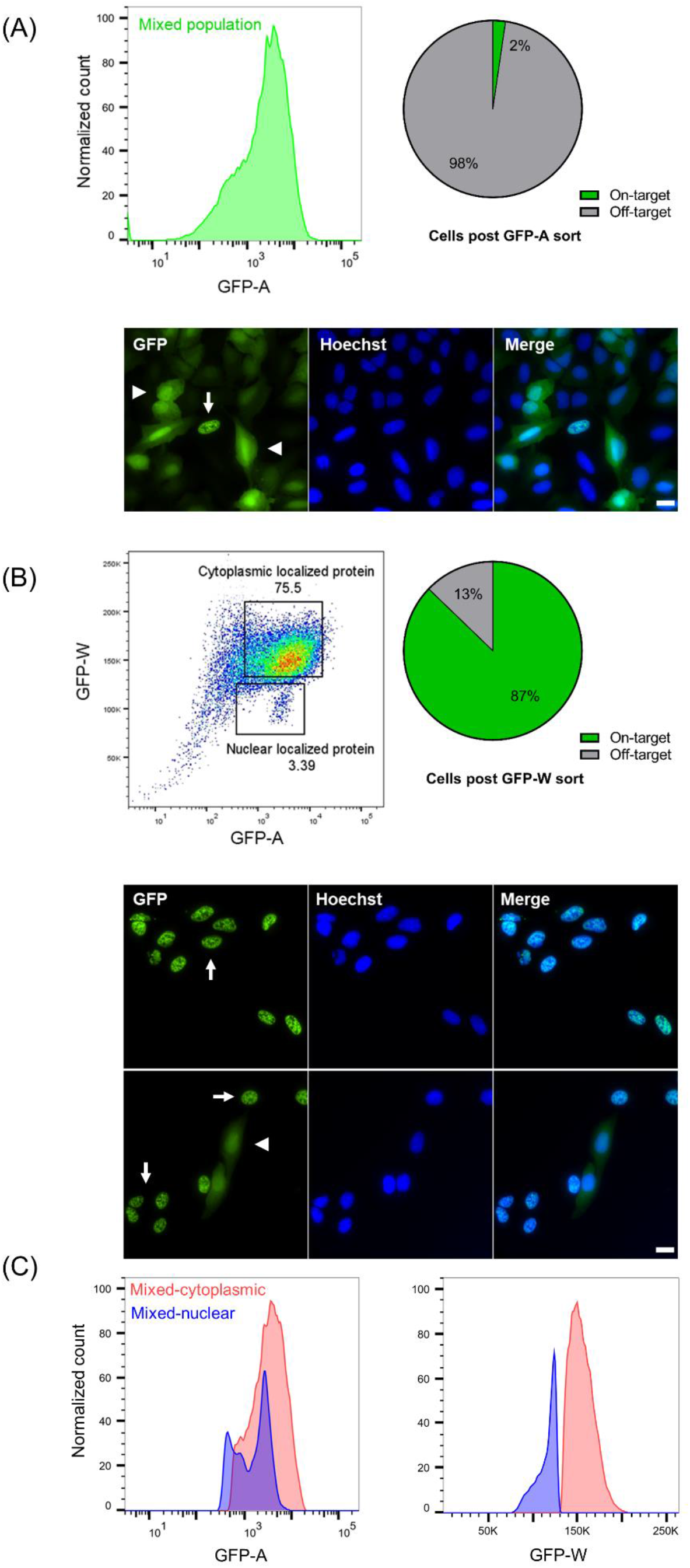
Utilization of new sorting gates for determining endogenous gene tagging. (A) GFP-A plot of cells after insert cassette transfection. Quantifications of positive cells show 2% of cells displaying on-target integration. Representative field shows a single positive cell (arrow), among many cells showing diffused green signal (arrowheads). Nuclear counterstain with Hoechst. Bars, 20 μm. (B) GFP-A versus GFP-W plot, showing two distinct populations with markedly different width of signal. After selecting for cells in the nuclear gate, quantifications of positive cells show significantly more (p<0.0001) cells displaying correct subcellular localization, reaching 87%. Representative fields show many green-positive cells (arrows), with a minority of cells showing diffuse green signal (arrowheads). (C) Graphs depicting GFP-A and GFP-W along an intensity axis, showing the mixed population split to the nuclear vs cytoplasmic localized gates. While the colonies overlap almost entirely in GFP-A, they show two distinct peaks along the GFP-W axis.

### 3.3 Single cell line generation after gating protocol

From the newly sorted nuclear localized population, single-cell lines were generated with relative ease, with over 80% of the cells examined after sorting showing the correct nuclear protein localization. This population was examined by flow cytometry as well. The cells were a cohesive population, with a large majority sitting firmly in the defined nuclear gate (Figure 3A). Cell line purity was validated by microscopy, and all cells showed on-target integration. Most cells expressed the protein in equal measure and displayed a uniform signal distribution when looking at GFP-A, with an extremely narrow peak because of their uniform expression (Figure 3B). Interestingly, when plotting this single-cell line against the histogram of the mixed cell population, we noticed that GFP-A overlaps entirely. This is in marked distinction to the GFP-W that yields two distinct peaks.

**FIGURE 3.**
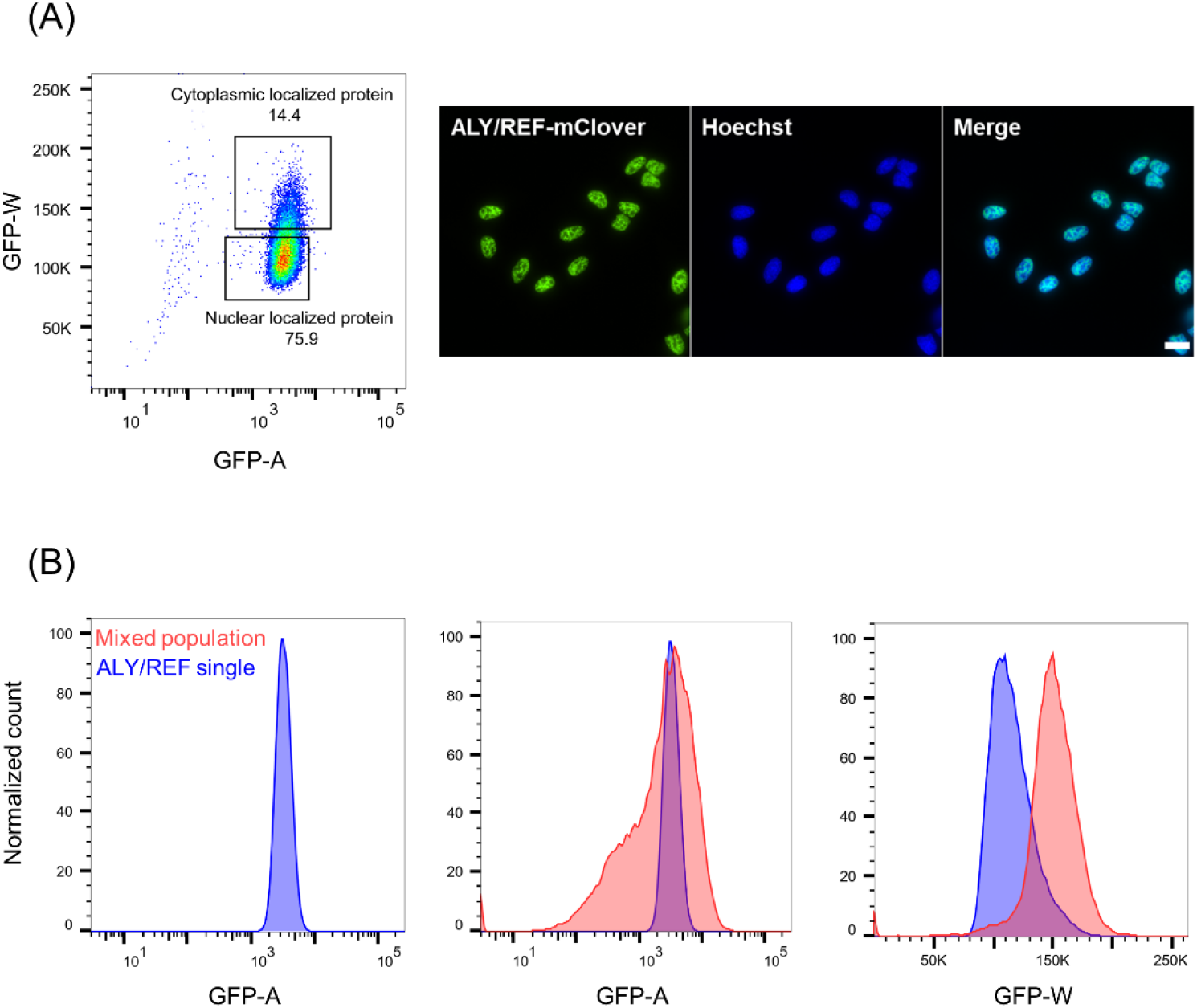
Single cell line generation after gating protocol. (A) GFP-A versus GFP-W plot, showing a cohesive cell population in the nuclear gate. Representative fields show only positive cells in the pure cell line. Nuclear counterstain with Hoechst. Bars, 20 μm. (B) Graphs depicting GFP-A and GFP-W along an intensity axis. The ALY/REF single cell line shows a narrow peak in GFP-A that overlaps entirely with the GFP-A of the mixed population of cells, but distinct GFP-W pattern.

## 4 DISCUSSION

Here we present a simple, high throughput, and easily reproducible method for enriching a cell population for positive on-target gene integration of a fluorescent tag into nuclear or cytoplasmic localized proteins. Our microscopic validation of the cytometry data demonstrates the quantitative component of this method as well. While standard sorting with flow cytometry accounts for overall signal, by utilizing the spatial component of expression it is possible to easily differentiate between cytoplasmic and nuclear signals. While this method can be easily implemented in a gene KI workflow, the principle of differentiating between cytoplasmic and nuclear signal can have broad applications.

It is usual to sort via fluorescent area into a 96-well plate to grow single cell colonies. While this can be effective, when integrating fluorescent proteins using a PCR cassette it is not uncommon for all wells to contain off-target green cells. Here, we dramatically accelerate cell line generation, such that close to 80% of the wells will contain a colony expressing the correct phenotype, and generate pure single cell lines with ease.

The single cell line we produced for the ALY/REF nuclear protein was validated using microscopy. It should be mentioned, while the population was uniform, there was always some bell curve of signal present whenever one acquires fluorescent data from a flow cytometer, it is therefore not surprising we see some signal spread into the cytoplasmic gate. We suspect that of the 14% of cells that appears in the cytoplasmic gate, some may be dividing cells, or simply a random error. A similar portion of cells can also be found in the cell line expressing nuclear localized protein which was used to create the gate.

The method described here can be easily applied to standard flow cytometers. We hope this simple yet powerfully effective tool will be used to generate endogenously tagged cell lines quickly and efficiently, in a way that was not previously realized.

## ACKNOWLEDGMENTS

We thank Jennifer I. C. Benichou (Bar Ilan University) for performing the statistical analysis.

## AUTHOR CONTRIBUTIONS

**Gabriel P. Faber:** Conceptualization (lead), data curation (lead), formal analysis (lead), methodology (equal), project administration (lead), validation (lead), visualization (lead), writing – original draft (lead), writing – review and editing (lead). **Hagit Hauschner:** Conceptualization (equal), data curation (equal), formal analysis (equal) methodology (lead), visualization (supporting), writing – review and editing (supporting). **Mohammad Khaled Atrash:** Data curation (supporting), methodology, (supporting), validation (supporting), visualization (equal), writing – review and editing (supporting). **Liat Bilinsky:** Conceptualization (supporting). **Yaron Shav-Tal:** Methodology (equal), project administration (equal), writing – original draft (equal), writing – review and editing (equal).

## REFERENCES

1. Dambournet, D., et al., Tagging endogenous loci for live-cell fluorescence imaging and molecule counting using ZFNs, TALENs, and Cas9. Methods Enzymol, 2014. 546: p. 139–60.

2. Jinek, M., et al., A programmable dual-RNA-guided DNA endonuclease in adaptive bacterial immunity. Science, 2012. 337(6096): p. 816–21.

3. Doudna, J.A. and E. Charpentier, Genome editing. The new frontier of genome engineering with CRISPR-Cas9. Science, 2014. 346(6213): p. 1258096.

4. Fueller, J., et al., CRISPR-Cas12a-assisted PCR tagging of mammalian genes. J Cell Biol, 2020. 219(6).

5. Scully, R., et al., DNA double-strand break repair-pathway choice in somatic mammalian cells. Nat Rev Mol Cell Biol, 2019. 20(11): p. 698–714.

6. Ramdzan, Y.M., et al., Tracking protein aggregation and mislocalization in cells with flow cytometry. Nat Methods, 2012. 9(5): p. 467–70.

7. Wienert, B., et al., Timed inhibition of CDC7 increases CRISPR-Cas9 mediated templated repair. Nat Commun, 2020. 11(1): p. 2109.

8. Hochberg-Laufer, H., et al., Uncoupling of nucleo-cytoplasmic RNA export and localization during stress. Nucleic Acids Res, 2019. 47(9): p. 4778–4797.

9. Faber, G.P., S. Nadav-Eliyahu, and Y. Shav-Tal, Nuclear speckles - a driving force in gene expression. J Cell Sci, 2022. 135(13).

10. Schwed-Gross, A., et al., Glucocorticoids enhance chemotherapy-driven stress granule assembly and impair granule dynamics, leading to cell death. J Cell Sci, 2022. 135(14).

11. Ashkenazy-Titelman, A., et al., RNA export through the nuclear pore complex is directional. Nat Commun, 2022. 13(1): p. 5881.

12. Gromadzka, A.M., et al., A short conserved motif in ALYREF directs cap- and EJC-dependent assembly of export complexes on spliced mRNAs. Nucleic Acids Res, 2016. 44(5): p. 2348–61.

13. Shi, M., et al., ALYREF mainly binds to the 5’ and the 3’ regions of the mRNA in vivo. Nucleic Acids Res, 2017. 45(16): p. 9640–9653.

